# Mitophagy is required for cell survival upon ultrasound and microbubble therapeutic exposure

**DOI:** 10.64898/2026.10.17.759693

**Authors:** Eleanor Cloves, Sahar A. Kamil, Nada Mayassi, Adelle Pinto, Raffi Karshafian, Costin N. Antonescu, Roberto J. Botelho

## Abstract

Ultrasound-stimulated microbubble (USMB) sonoporation treatment is an emerging strategy for cancer therapy that enhances drug delivery by perforating the plasma membrane. This membrane permeabilization also causes cellular metabolic stress as cells leak metabolites such as ATP and amino acids. Collectively, cells may die in response to excess membrane damage and stress. However, many cells survive the USMB insult by engaging membrane repair and adaptation pathways such as stimulation of AMPK. In this study, we investigated the impact of USMB treatment on mitochondria and their role in post-exposure cellular survival. We found that USMB causes mitochondrial fragmentation and decreases mitochondrial membrane potential. Subsequently, autophagy and Parkin-dependent mitophagy are triggered post-USMB treatment in an AMPK-dependent mechanism. Importantly, Parkin silencing and PINK1 inhibition augment cell death when combined with USMB exposure, suggesting that mitophagy is an important cell survival process in response to USMB. We propose that combining USMB with inhibitors of mitophagy and metabolic stress adaptation is a strong candidate for therapeutic development to treat solid tumours.

## Introduction

Ultrasound-stimulated microbubble (USMB) sonoporation is an emerging strategy for precision killing of tumour cells (Czarnota et al., 2012; Dasgupta et al., 2023; van Wamel et al., 2004). During USMB sonoporation, microbubbles within the ultrasound field undergo both stable and inertial cavitation, generating fluid microjets and shear forces that disrupt the plasma membrane of proximal cells, causing bidirectional diffusion into and out of the cell (Cattaneo et al., 2025; Okada et al., 2005; van Wamel et al., 2006). This enhances the uptake of impermeable extracellular drugs while the cell significantly loses proteins and metabolites (Hussein et al., 2017; Lo et al., 2023; Meijering et al., 2009; van Wamel et al., 2004). Cellular responses to USMB are heterogeneous, such that a proportion of cells die while others survive under the same ultrasound exposure conditions. The extent of cell death depends on parameters such as ultrasound frequency, acoustic pressure, duty cycle, microbubble concentration, and the cell model. For example, studies using i*n vitro* cancer cell models exposed under commonly used USMB conditions have reported approximately 30-50% cell death, leaving a substantial surviving population (Almasri and Karshafian, 2022; Karshafian et al., 2009). Nevertheless, preclinical studies have demonstrated that USMB sonoporation, especially when combined with other therapeutic strategies such as radiation and chemotherapeutics, suppresses tumour growth and enhances tumour cell death (Czarnota et al., 2012; Goertz et al., 2012; Liu et al., 2024; Sharma et al., 2022).

A key question is how cells survive and adapt to the initial USMB sonoporation insult. For example, cells that survived USMB sonoporation not only engaged in membrane repair processes, but also underwent substantial metabolic stress because of the loss of proteins, ATP, amino acids, and glycolytic metabolites (Hussein et al., 2017; Lo et al., 2023). As a result, we previously observed that survivor cells activate Adenosine Monophosphate (AMP)-activated protein kinase (AMPK), a central metabolic regulator that promotes ATP recovery (Lo et al., 2023). In fact, using pharmacological and shRNA-mediated silencing of AMPKα1/α2, we demonstrated that AMPK is required for ATP recovery in surviving cells. Moreover, arresting AMPK activity enhanced USMB-mediated cell death, highlighting the therapeutic potential of combining USMB sonoporation with blockers of AMPK or other metabolic adaptation pathways (Lo et al., 2023).

Mitochondria are critical organelles in metabolic adaptation, responsible for oxidative phosphorylation and ATP synthesis, citric acid cycle, fatty acid β-oxidation, amino acid shunts, and more (Birsoy et al., 2015; Nunnari and Suomalainen, 2012; Spinelli and Haigis, 2018). Mitochondria are highly dynamic organelles that undergo fission-fusion cycles to maintain their genetic and metabolic quality, such as membrane potential and oxidative state (Adebayo et al., 2021; Ng et al., 2021). For example, mitochondrial fragmentation is associated with loss of membrane potential, isolation of damaged mitochondrial components for degradation, and, under severe stress, can even induce apoptosis (Suen et al., 2008; Twig and Shirihai, 2011). Given their centrality, there are multiple mitochondrial quality control systems that repair or eliminate permanently damaged mitochondria (Li et al., 2025).

Mitophagy is the selective elimination of mitochondria via autophagy either due to damage, aging, or as a scavenging tool during nutrient depletion (Lu et al., 2023). The PINK1-Parkin mitophagy pathway is arguably the best understood (Pickrell and Youle, 2015). Loss of mitochondrial membrane potential and other cues lead to stabilization of the kinase PINK1 on the mitochondrial surface; PINK1 phosphorylates ubiquitin associated with mitochondrial outer membrane proteins and recruits the E3 ubiquitin ligase Parkin, which is also phosphorylated by PINK1 (Lazarou et al., 2015; Narendra et al., 2010; Vives-Bauza et al., 2010). Parkin then ubiquitylates mitochondrial outer membrane proteins, which subsequently leads to recruitment of autophagy receptors to initiate autophagic enclosure of the depolarized mitochondrion (Geisler et al., 2010; Heo et al., 2015; Lazarou et al., 2015; Narendra et al., 2008, 2010; Vives-Bauza et al., 2010). Mitophagosomes then fuse with lysosomes to degrade and recycle the components – this is referred to as mitophagic flux and can be assessed by acidification of mitochondrial-targeted Keima (Sun et al., 2015). Overall, mitophagy is critical for cells to maintain their mitochondrial quality, or, as with bulk autophagy, to scavenge raw materials during nutrient deprivation conditions. This may be particularly true in metabolically stressed tumour cells, common in triple-negative breast cancers, which exhibit metabolic plasticity in both hypoxic and nutrient-deprived microenvironments (Wang et al., 2020).

Given the metabolic stress imposed on cells during USMB and the role of AMPK in resisting death, we hypothesized that mitochondrial dynamics would play a central role in cell adaptation and survival in response to USMB. In fact, we first observed that USMB caused extensive mitochondrial fragmentation, loss of membrane potential, and boosted oxidative stress. We further observed Parkin-dependent mitophagy and mitophagic flux post-USMB. Importantly, inhibition of Parkin or PINK1 during USMB augmented cell death compared to individual treatments. Collectively, our findings identify mitophagy as an important adaptive response that enables cell survival following USMB exposure and position this axis as a potential therapeutic vulnerability to treat tumours like triple-negative breast cancers.

## Results

### USMB alters mitochondrial morphology and motility

We previously observed that cells lose intracellular proteins, ATP, and other metabolites after USMB, and require AMPK to recover and survive post-USMB (Hussein et al., 2017; Lo et al., 2023). Since mitochondria play a central role in metabolite and ATP generation, we then inquired about the role of mitochondria in cell survival post-USMB treatment. To begin addressing this, we used a strategy depicted in Fig. 1A to examine if USMB treatment altered mitochondrial dynamics, including their morphology and motility, in MDA-MB-231 triple-negative breast cancer cells labelled with MitoTracker Green before and after USMB exposure.

**Figure 1.**
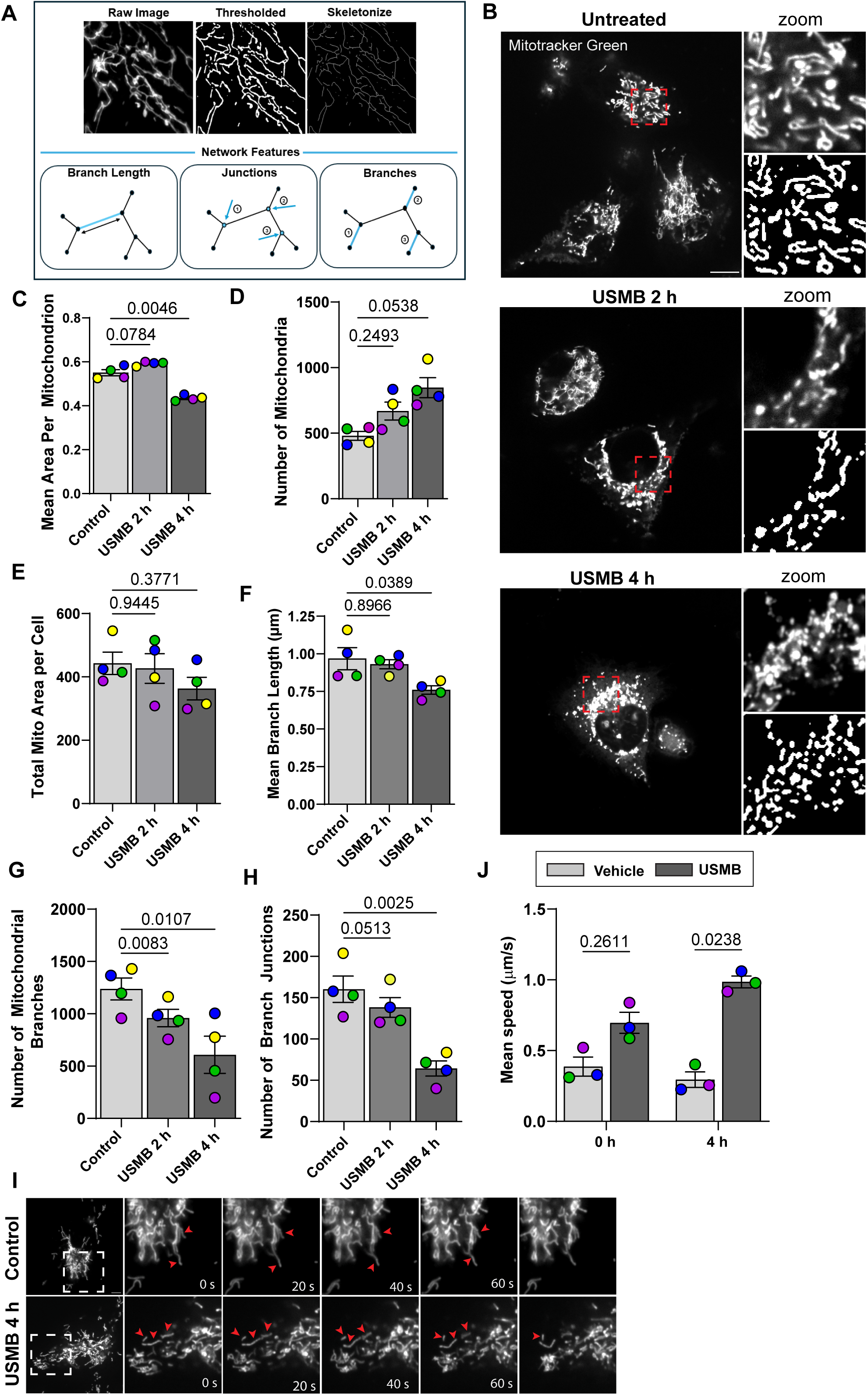
Increased mitochondrial fragmentation and motility in MDA-MB-231 cells after USMB sonoporation. **A.** Schematic illustrating the image-analysis workflow used to quantify mitochondrial network parameters using segmented images. **B.** Mitochondria in MDA-MB-231 cells labelled with MitoTracker Green before USMB sonoporation (control), or 2 or 4 h after USMB. Enlarged views of the boxed regions and corresponding segmented mitochondrial outputs are shown to the right. **C**-**H.** Quantification of mitochondrial morphological parameters including mean area per mitochondrion (C), number of mitochondria (D), total area of mitochondria (E), mitochondrial branch length (F), number of branches (G), and branch junctions (H) in MDA-MB-231 cells before, or 2 or 4 h post-USMB sonoporation. Data are shown as mean ± SEM from N=4 independent replicates, where 100-200 cells per condition per experiment were quantified. **I.** Mitochondria in MDA-MB-231 cells labeled with MitoTracker Deep Red, imaged using time-lapse spinning disk microscopy, showing motility before USMB treatment (control) or 4 h post-USMB. Dashed boxes indicate regions shown at higher magnification, and arrowheads indicate examples of moving mitochondrial structures. These data are supported by corresponding Videos S1 and S2. **J.** Quantification of the mean speed (µm/s) of the mitochondria at 0 and 4 h, with and without USMB treatment (N=3). Data in C-H were statistically compared by one-way ANOVA, followed by Dunnett’s Multiple Comparisons test and p values are as shown. Data in J were analysed by two-way ANOVA and Šidak’s multiple comparison post hoc test. Scale bar = 5 µm.

In untreated cells (prior to USMB exposure), mitochondria were typically elongated and branched, as expected (Fig. 1B). However, USMB induced major changes in the mitochondrial network, including a robust reduction in mean area per mitochondrion, while the mitochondrial count per cell increased. In addition, at 4 h post-USMB, mitochondria showed reductions in branch and junction numbers. Despite these changes, total mitochondrial area per cell remained relatively stable (Fig. 1B-H). Importantly, MDA-MB-468 cells underwent similar reductions to mitochondrial size, branch length, and network complexity in response to USMB, indicating that these observations were not specific to MDA-MB-231 cells (Sup. Fig. S1). To determine if the mitochondrial remodeling was sustained at later timepoints, mitochondrial morphology was examined at 24 and 48 h post-USMB. At these later timepoints, mean mitochondrial branch length and mitochondrial branch number were no longer significantly altered, suggesting at least partial recovery of mitochondrial morphology occurring after the acute 4 h response (Sup. Fig. S2). Together, these findings indicate that USMB-induced mitochondrial fragmentation occurs acutely at 4 h, while aspects of mitochondrial morphology largely recover after 24 h. Aside from the changes to mitochondrial morphological properties, mean mitochondrial speed was increased in cells 4 h post-USMB relative to control cells (Fig. 1I, J; Sup. Video S1, S2). Overall, our observations indicate that USMB triggers fundamental changes to mitochondrial dynamics by promoting fragmentation and increased motility.

### USMB triggers mitochondrial depolarization and oxidative stress

Given the substantial remodeling of the mitochondrial network organization following USMB, we next examined whether USMB also impacted mitochondrial membrane potential (ΔΨm) and cellular oxidative stress. To assess the mitochondrial membrane potential, MDA-MB-231 cells were stained with TMRE alongside MitoTracker Green to visualize total mitochondrial mass under control and 4 h conditions following USMB treatment. Relative to control, there was a significant, albeit small, reduction in the TMRE signal relative to MitoTracker Green 4 h post-USMB treatment, suggesting a partial loss of mitochondrial membrane potential (Fig. 2A, B).

**Figure 2.**
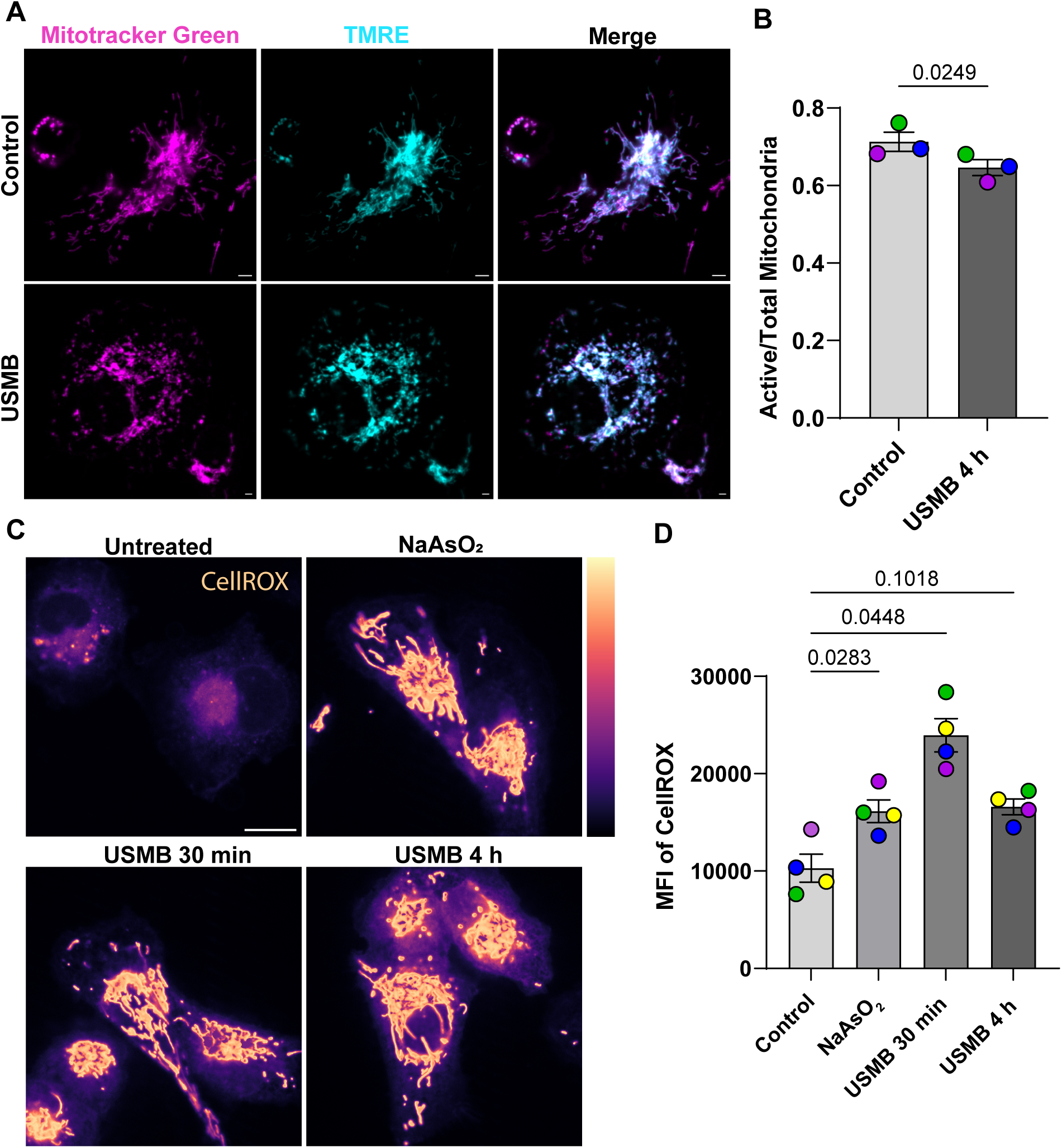
USMB disturbs mitochondrial membrane potential and induces oxidative stress. **A.** Spinning disk microscopy images of MDA-MB-231 cells stained with MitoTracker Green and TMRE before or 4 h following USMB treatment. Merged images are shown at the right. **B.** Quantification of the ratio of TMRE-positive (active) mitochondria to total mitochondrial area (MitoTracker Green-positive). Data are shown as normalized mean ± SEM from N=3 independent replicates, where 50-100 cells per condition per experiment were quantified. **C.** Spinning disk confocal microscopy images of CellROX-stained MDA-MB-231 cells before, 30 min, or 4 h post-USMB; NaAsO_2_ is a positive control for oxidative stress. **D.** Quantification of CellROX mean fluorescence intensity in control, NaAsO_2_, 30 min USMB, and 4h USMB. Data are shown as normalized mean ± SEM from N=4 independent replicates, where 75-125 cells per condition per experiment were quantified. Data in B were statistically compared with a student’s paired t-test. Data in D were compared by one-way ANOVA, followed by Dunnett’s Multiple Comparisons post-hoc test; p values are shown. In A, Scale bar = 1 µm. In C, scale bar = 5 µm.

Given the close relationship between mitochondrial membrane potential and ROS production during mitochondrial dysfunction (Bulthuis et al., 2019; Kuznetsov et al., 2017), we next examined whether USMB treatment altered oxidative stress levels by staining with CellROX Deep Red. Relative to resting conditions, the CellRox signal intensity was significantly higher in cells 30 min post-USMB exposure compared to untreated cells. By 4 h post-USMB, the CellROX fluorescence remained elevated relative to control cells, although this difference was no longer statistically significant (Fig. 2C, D). Interestingly, the early exposure to USMB produced greater oxidative stress than cells incubated with sodium arsenite, a known ROS inducer (Fig. 2C, D). Collectively, these results demonstrate that USMB hinders mitochondrial membrane potential and promotes oxidative stress, suggesting that mitochondrial dysfunction occurs following USMB treatment.

### USMB triggers autophagy and Parkin recruitment to mitochondria

Mitochondrial depolarization and increased oxidative stress following USMB suggested that mitochondria become functionally compromised and may promote turnover. Autophagy is a degradative pathway that removes cytoplasmic material through lysosomal degradation. Mitophagy is the selective autophagic clearance of mitochondria, particularly those that are damaged or dysfunctional (Li et al., 2015; Song et al., 2018). To assess whether USMB induced autophagic responses, we examined LC3-positive puncta formation in MDA-MB-231 cells. LC3-II, the membrane associated form, is recruited to structures representing all stages of autophagy including phagophores, autophagosomes and autolysosomes, making LC3 puncta a marker of autophagic structures (Kabeya et al., 2000).

In untreated cells, LC3^+^ puncta were sparse, consistent with the expected low basal autophagy. However, USMB-treated cells exhibited a significant increase in LC3^+^ puncta, particularly at 4 h following USMB treatment (Fig. 3A, B). These findings suggest that USMB stimulates autophagy. The fraction of mitochondrial signal overlapping with LC3-positive structures was also increased following USMB treatment (Fig. 3C) suggesting enhanced association of autophagic structures with mitochondria.

**Figure 3.**
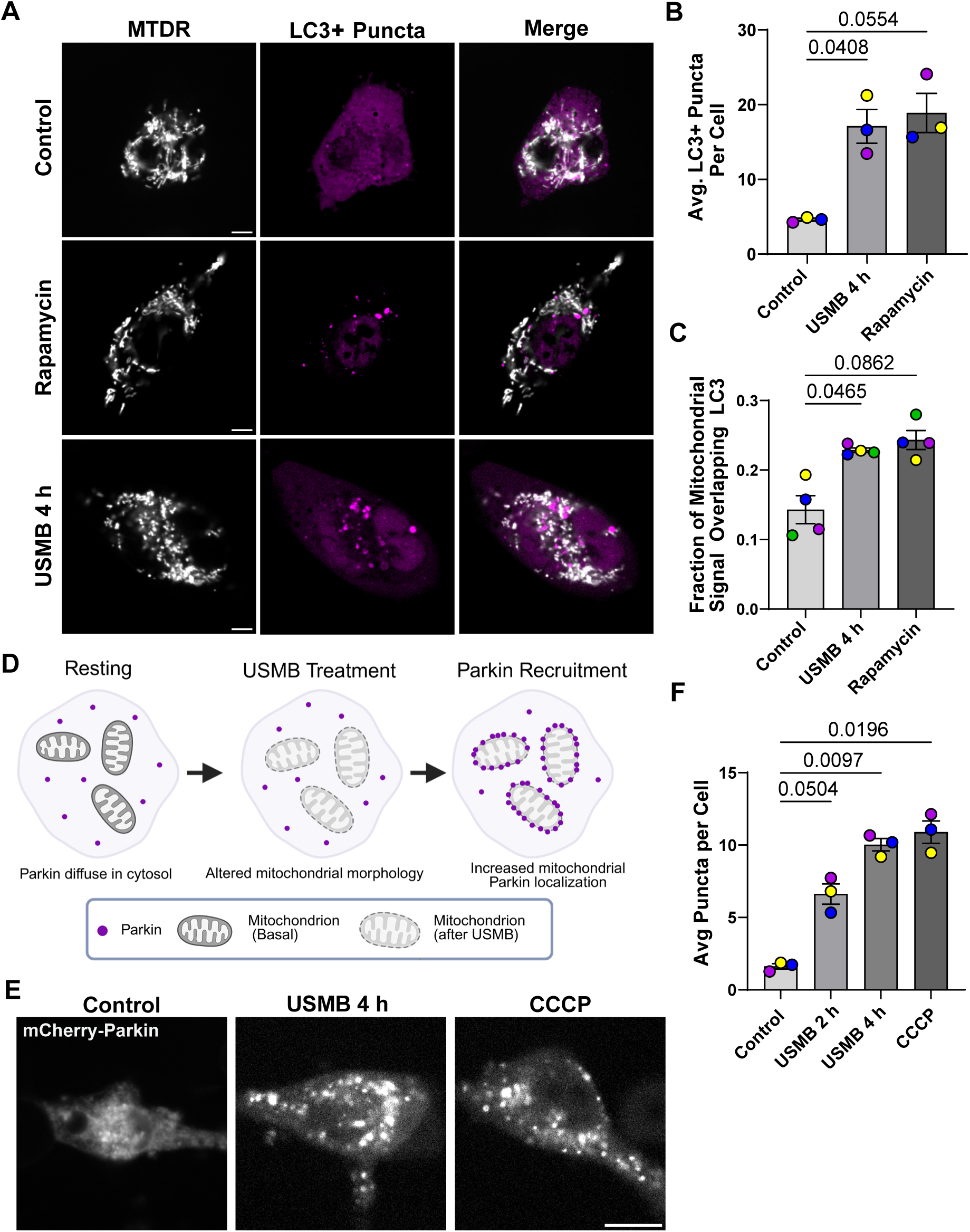
USMB sonoporation triggers autophagy and Parkin recruitment to mitochondria. **A**. Representative micrographs of MDA-MB-231 cells stably expressing tandem mCherry-eGFP-LC3 reporter and labelled with MitoTracker Deep Red (MTDR) under control conditions, or after exposure to 1 µM rapamycin, or 4 h USMB. The mCherry channel was used to visualize total LC3-positive puncta. Merged images demonstrate the spatial relationship between LC3-positive structures and mitochondria. **B.** Quantification of the average number of LC3-positive puncta per cell under control, 4h USMB, or rapamycin conditions. Data are shown as mean ± SEM from N=3 independent experiments, with approximately 50-100 cells analyzed per condition per experiment. **C.** Quantification of the fraction of mitochondrial signal overlapping with LC3-positive puncta using the Manders’ coefficient. **D.** Schematic illustrating the recruitment of Parkin following USMB treatment. Under resting conditions, Parkin is diffuse within the cytosol. Following USMB treatment, Parkin forms puncta that show increased association with mitochondria. **E.** Representative micrographs of MDA-MB-231 cells transiently expressing mCherry-Parkin under control conditions, or 4 h following USMB treatment, or following treatment with CCCP. **F.** Quantification of the average number of mCherry-Parkin puncta per cell under control, 2 h or 4 h following USMB treatment, or with CCCP. Data in B, C and F were analyzed by one-way ANOVA followed by Dunnett’s multiple comparisons test. Data are presented as mean ± SEM, from N= 3 and 4, respectively. Scale bar = 5 µm.

Detection of LC3 puncta could reflect induction of bulk autophagy or the selective autophagic degradation of mitochondria by mitophagy. Given the mitochondrial alterations observed earlier, we next investigated recruitment of the E3 ubiquitin ligase Parkin to mitochondria, which is selective for mitophagy and not seen in bulk autophagy. We transiently expressed mCherry-Parkin in MDA-MB-231 cells to monitor its recruitment to mitochondria following USMB. Under untreated conditions, Parkin displayed a diffuse cytosolic localization. In contrast, cells treated with USMB exhibited an accumulation of Parkin puncta, similar to cells treated with the positive control for mitophagy, CCCP (Fig. 3D-F). Collectively, these findings indicate that USMB activates an autophagic response associated with mitochondria and promotes Parkin recruitment following mitochondrial stress.

### Autophagic and mitophagic flux proceeds after USMB treatment

USMB induced LC3-positive puncta and Parkin recruitment to the mitochondria. However, the completion of autophagy requires fusion of autophagosomes with lysosomes, culminating in the subsequent degradation of cargo, largely referred to as autophagic flux (Zhang et al., 2013). We proceeded to examine whether LC3-positive structures and mitochondrial cargo progressed into acidic compartments following USMB treatment.

To assess autophagic flux, we utilized MDA-MB-231 cells expressing the mCherry-GFP-LC3 tandem reporter which distinguishes non-acidified autophagosomes (mCherry^+^/GFP^+^) from acidified autolysosomes (mCherry^+^/GFP^-^) since the GFP fluorophore is quenched in acidic environments (Kimura et al., 2007). In control cells, acidified LC3-positive puncta were relatively sparse, indicating a low steady-state abundance of LC3-positive autolysosomal structures. Following USMB treatment, cells displayed an increased proportion of acidified LC3-positive puncta relative to total LC3-positive puncta, consistent with a rapid progression of LC3-positive structures into acidic autolysosomal compartments (Fig. 4A, B).

**Figure 4.**
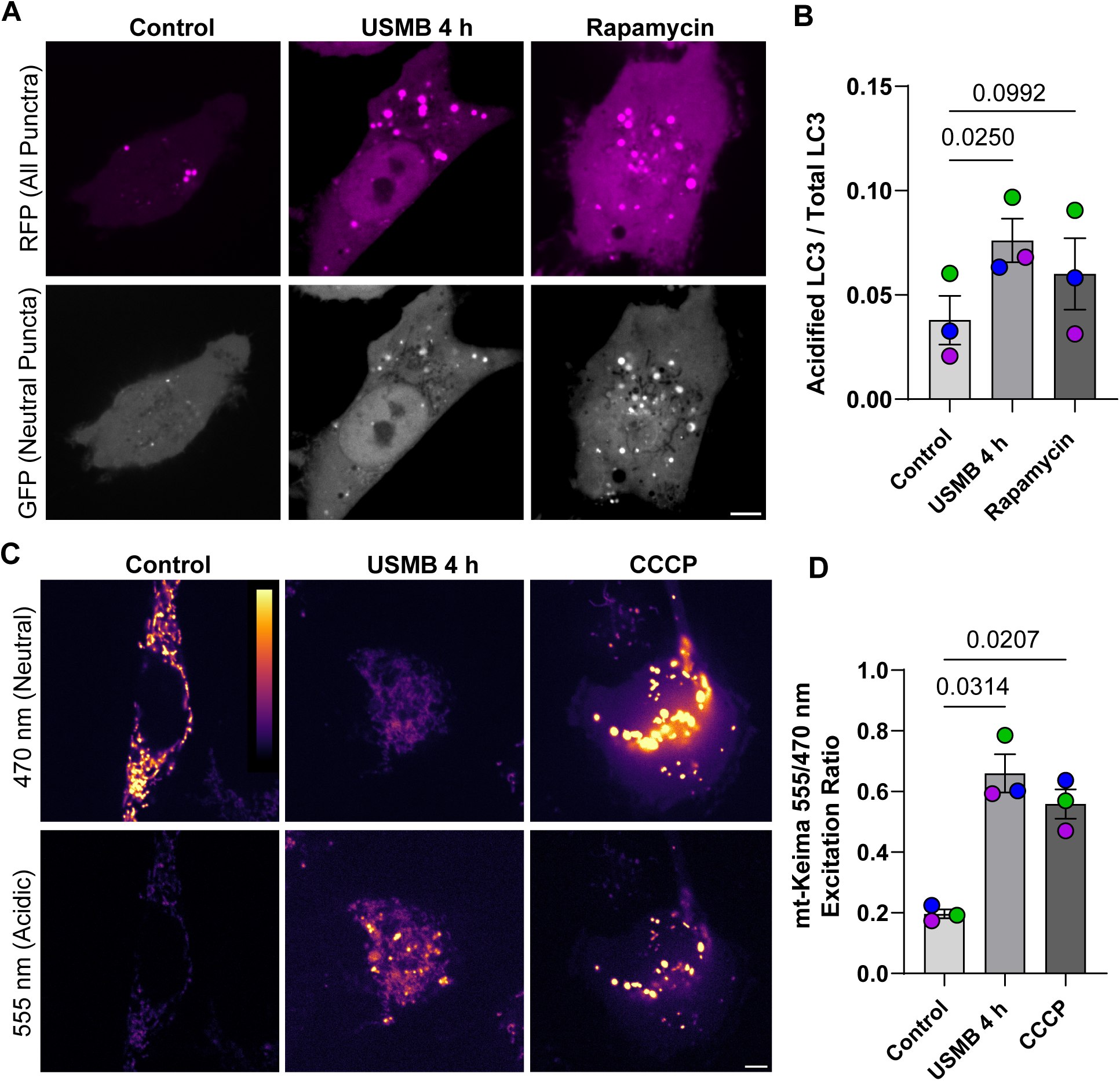
USMB induces autophagosome accumulation and promotes mitochondrial delivery to acidic compartments. **A.** Representative spinning disk microscopy images of MDA-MB-231 cells stably expressing mCherry-eGFP-LC3. mCherry labels total LC3-positive puncta, whereas GFP signal is quenched in acidic compartments, allowing for the determination of acidified vs non-acidified LC3^+^ puncta. **B.** Quantification of GFP fluorescence intensity within an mCherry-defined LC3-positive mask, reflecting acidification of LC3-positive compartments. Data are shown as normalized mean ± SEM from N=3 independent replicates, where 50-100 cells per condition per experiment were quantified. **C.** Representative micrographs of mt-Keima fluorescence after excitation at 470 nm (preferential excitation under neutral conditions) or 555 nm (preferential excitation under acidic conditions) under control conditions, or 4h following USMB treatment, or CCCP. Display is in false-colour using the LUT mpl-inferno. **D.** Quantification of the mt-Keima fluorescence ratio in control cells, or after USMB or CCCP treatments (N = 3). Data in B and D were statistically compared by one-way ANOVA, followed by Dunnett’s Multiple Comparisons test and p values are as shown. Scale bar = 5 µm.

To determine if mitochondria were delivered to acidified compartments following USMB treatment, we used the mitochondrial-targeted pH-sensitive reporter mt-Keima. Mt-Keima is targeted to the mitochondrial matrix, which allows for changes to its excitation profile to report exposure of mitochondrial material to an acidic environment rather than changes in proton accumulation within the intermembrane space associated with mitochondrial membrane potential (Katayama et al., 2011; Sun et al., 2015). Control cells exhibited a relatively low 555/470 excitation ratio, consistent with most mitochondria residing in a neutral environment. In contrast, USMB treatment significantly increased the mt-Keima 555/470 excitation ratio at 4 h, indicating increased localization of mitochondrial material within acidic compartments (Fig. 4C, D). This was like the increase observed following treatment with the mitochondrial depolarizing agent CCCP. As a corollary, these data show that USMB promotes the delivery of mitochondria into acidic compartments, consistent with active mitophagic flux.

### USMB-induced mitophagic signaling is partially dependent on AMPK activity

To investigate the signalling mechanisms that contribute to the mitophagic response following USMB treatment, we next examined the role of AMPK. MDA-MB-231 cells transiently expressing mCherry-Parkin were treated with vehicle or the AMPK inhibitors Compound C (dorsomorphin) or BAY-3827 prior to USMB treatment. Consistent with our previous observations, USMB treatment promoted the relocation of Parkin from the cytosol to discrete punctate structures associated with mitochondria in vehicle-treated cells (Fig. 5A, B). In comparison, Compound C- or BAY-3827 significantly blunted the number of Parkin puncta formed after USMB (Fig. 5A, B; compare USMB + vehicle vs. USMB + AMPK inhibitors), albeit this was a partial arrest (Fig. 5A, B; compare no USMB vs. USMB for each of the AMPK inhibitors). Nevertheless, these data indicate that AMPK activity is required to fully promote mitophagy triggered by USMB treatment.

**Figure 5.**
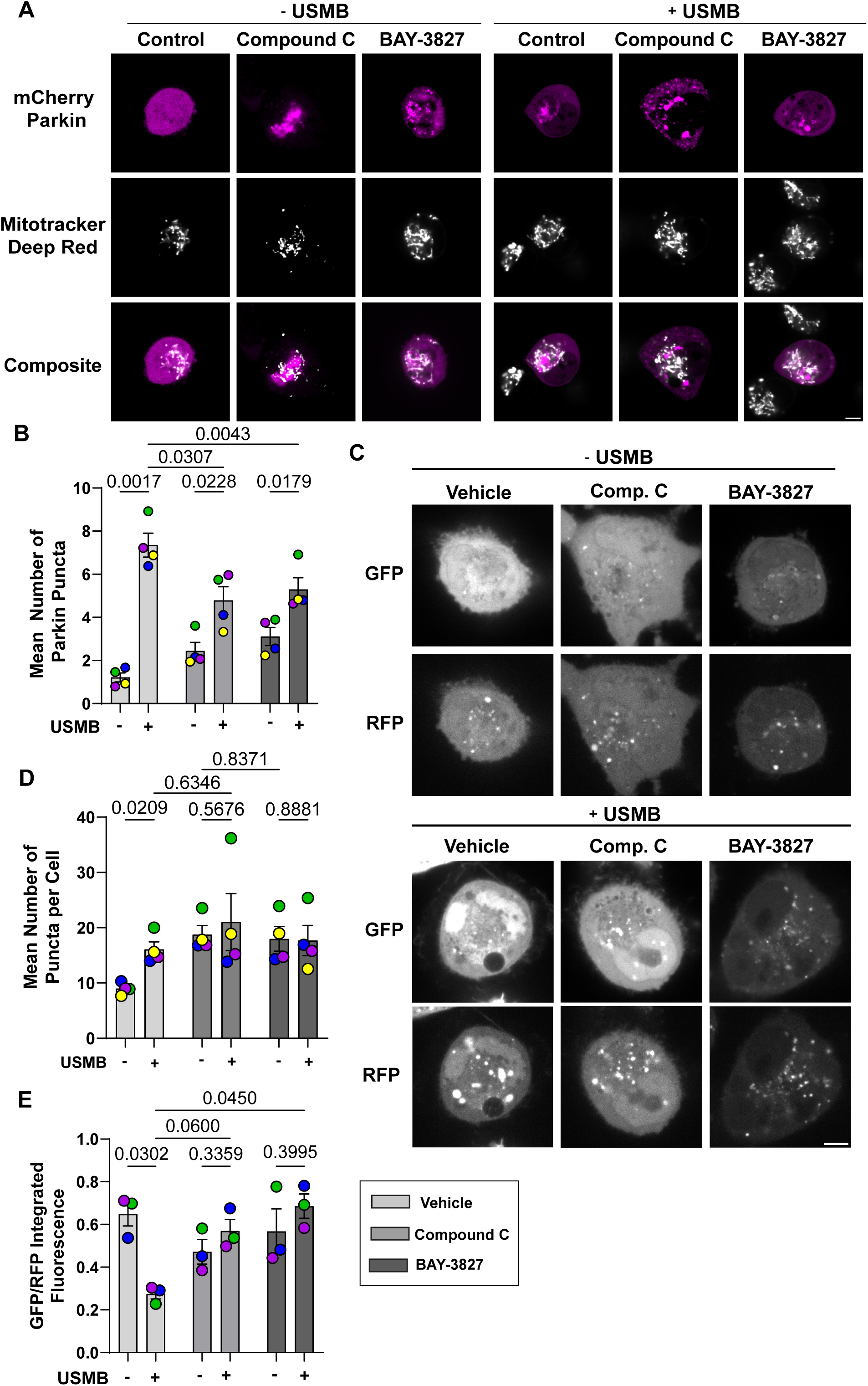
USMB-induced mitophagy is dependent on AMPK activity. **A, C.** Representative micrographs of MDA-MB-231 cells transiently expressing mCherry-Parkin and stained with MitoTracker Deep Red (A) or expressing mCherry-eGFP-LC3 (C) before or after USMB and pre-treated with vehicle or AMPK inhibitors. Scale bar = 5 µm. **B, D** and **E.** Quantification of mean number of Parkin puncta (B), of the mean number of LC3+ puncta per cell (D), and of GFP/RFP integrated fluorescence of LC3 puncta (E) in cells pre-treated with vehicle, or the AMPK inhibitors Compound C and BAY3827, with and without USMB treatment. Statistical comparison was performed as a two-way ANOVA followed by Tukey’s multiple comparison post hoc test for B and E, while a two-way ANOVA followed by Šidak’s multiple comparison post hoc test was used for D. Data are shown as mean ± SEM with N = 4 (B, D) and N = 3 (E) independent replicates. P-values are as shown.

To determine whether AMPK activity also contributed to the autophagic response following USMB, we next examined the tandem GFP-RFP-LC3 reporter cells treated with Compound C or BAY-3827 (Fig. 5C-E). Once again, USMB boosted LC3-positive puncta (Fig. 5C, D) and rapid acidification of autophagosomes (Fig. 5C, E). However, when cells were treated with Compound C or BAY-3827 to impair AMPK activity, this augmented the number of LC3 puncta on their own (Fig. 5C, D). We could not detect an additive effect when combining AMPK antagonists with USMB (Fig. 5C, D) precluding our ability to determine if AMPK activity is required for USMB-mediated increase in LC3-labelled autophagosomes. We suspect that the increase in LC3 puncta in cells arrested for AMPK activity may be due to an obstruction in autophagosome turnover or flux since Compound C and BAY-3827 maintained high GFP/RFP ratio indicating low acidification of autophagosomes (Fig. 5C, E); this is consistent with past observations that activated AMPK promotes V-ATPase activity and autophagosome acidification (McGuire and Forgac, 2018).

### Blocking the PINK1-Parkin pathway enhances cell death following USMB

Since we had established that USMB treatment promotes mitophagy, we next asked whether this phenomenon contributes to cellular survival following USMB-induced stress. To first test this by gene silencing, Parkin was silenced using two independent siRNA oligonucleotides. Their knockdown efficiency was assessed in HeLa cells overexpressing Parkin, demonstrating reduced Parkin protein abundance relative to the non-targeting siRNA control (Sup. Fig. 3A, B). Following Parkin depletion, cell growth and death were monitored using Incucyte live-cell imaging. Parkin silencing alone produced a modest effect on cell growth and death compared with non-targeting control conditions. In contrast, combining Parkin silencing with USMB treatment reduced cell growth (Fig. 6A, B, Sup. Videos S3-S8) and increased death compared with non-targeting control conditions (Fig. 6A, C; Sup. Video S3 through S8). To account for differences in the cell number of the remaining cell population, cell death was additionally quantified relative to cell confluence. This demonstrated a clearly enhanced cell death level relative to confluency following combined Parkin depletion and USMB treatment (Fig. 6D). A similar sensitization to death phenotype was also observed in the MDA-MB-468 cell line (Sup. Fig. 3C-F). Together, these findings indicate that Parkin contributes to cell survival following USMB-induced stress.

**Figure 6.**
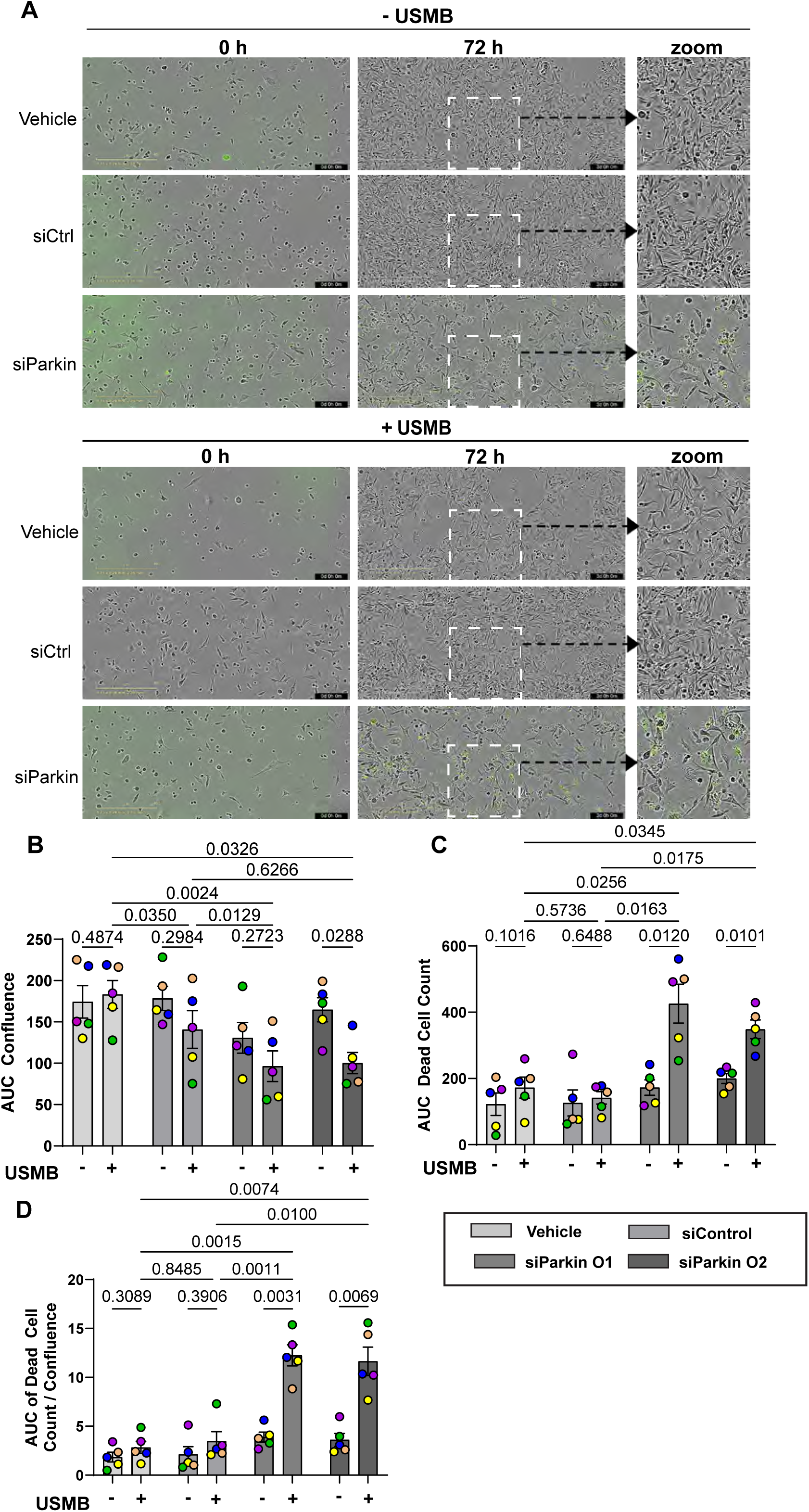
Parkin silencing sensitizes cells to USMB-induced cell death. **A.** Representative IncuCyte merged phase and fluorescence images acquired over 72 h showing CellTOX Green positive cells under vehicle, non-targeting siRNA control, or silenced Parkin conditions, in the presence or absence of USMB. **B.** Quantification of cell confluence measured hourly over 72 h, expressed as the area under the curve (AUC) of cell confluence for vehicle, siControl-transfected cells, and cells transfected with two independent Parkin-targeting siRNAs (siParkin O1 and siParkin O2), with or without USMB treatment. **C.** Quantification of cumulative cell death over 72 h, expressed as the AUC of CellTOX-positive dead-cell count. **D.** Quantification of cell death relative to cell abundance, expressed as the AUC of dead-cell count normalized to cell confluence. Data are shown as mean ± SEM from N=5 independent experiments. Statistical comparisons were performed using a 2-way ANOVA, followed by Šidak’s multiple comparisons test. P-values are as shown.

Having established this relationship by gene silencing, we next asked whether pharmacological disruption of the PINK1-Parkin pathway could similarly sensitize cells to USMB. To test this, we used BIIB-057, a compound recently reported to inhibit PINK1 but that has also been shown to target SYK kinase (Rasool et al., 2024). While MDA-MB-231 cells do not appear to express Syk (Coopman et al., 2000; Mahabeleshwar and Kundu, 2003; Wang et al., 2003), we nevertheless compared BIIB-057 with Entospletinib, a selective Syk inhibitor to distinguish between these two activities (Currie et al., 2014). Notably, Entospletinib did not increase death relative to the corresponding vehicle control (Fig. 7A-C). In comparison, BIIB-057 alone had a modest effect on cell growth and death (Fig. 7A-C). However, when we normalized cell death to confluency to account for differences in cell number, we observed a statistically significant increase in cell death in cells exposed to the combined treatment of BIIB-057 and USMB relative to other conditions (Fig. 7D). These findings suggest that pharmacological inhibition of PINK1 sensitizes cells to USMB-induced stress and support a protective role for the PINK1-Parkin mitochondrial pathway. These findings are consistent with the Parkin-silencing experiments and further support a protective role for the PINK1-Parkin pathway following USMB-induced stress. Collectively, these observations suggest that disruption of this mitochondrial quality control response may enhance the cytotoxic effects of USMB.

**Figure 7.**
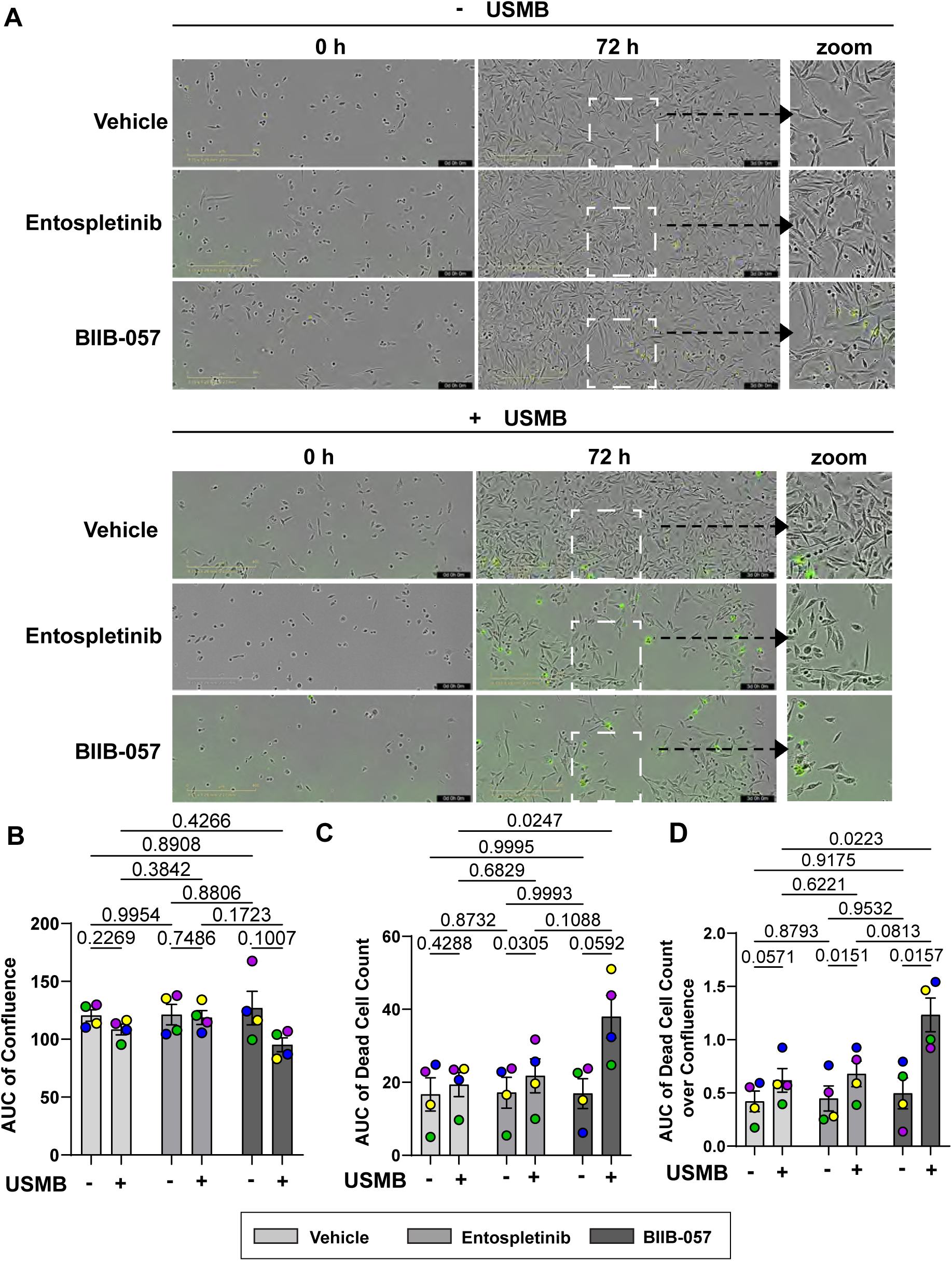
Pharmacological inhibition of PINK1 sensitizes cells to USMB-induced cell death. **A.** Representative IncuCyte phase-contrast and CellTOX Green fluorescence images acquired at 0 and 72 h from MDA-MB-231 cells treated with vehicle, the selective SYK inhibitor entospletinib, or the PINK1 inhibitor BIIB-057, with or without USMB treatment. Enlarged images are shown at the right. Scale bar = 400 µm. **B.** Quantification of cell confluence measured hourly over 72 h, expressed as the area under the curve (AUC) of cell confluence for vehicle, entospletinib or BIIB-057, with or without USMB treatment. **C.** Quantification of cumulative cell death over 72 h, expressed as the AUC of CellTOX-positive dead-cell count. **D.** Quantification of cell death relative to cell abundance, expressed as the AUC of dead-cell count normalized to cell confluence. Data are shown as mean ± SEM from N=4 independent experiments where 150-200 cells per condition per experiment were quantified. Statistical comparisons were performed using a 2-way ANOVA, followed by Tukey’s post hoc test. P values are as displayed.

## Discussion

USMB treatment may enable precision killing of tumour cells within an ultrasound field by compromising the plasma membrane and causing bidirectional free diffusion of extracellular and cytoplasmic molecules (Hussein et al., 2017; Lentacker et al., 2014; Lo et al., 2023; Ohl et al., 2006; Okada et al., 2005; van Wamel et al., 2006, 2004). However, a proportion of cells exposed to USMB survive because of insufficient damage and/or engagement of membrane repair and stress-adaptation pathways (Duan et al., 2021; Lee et al., 2025; Leow et al., 2015; Okada et al., 2005). In fact, we demonstrated that USMB sonoporation promoted AMPK activity, which helped recover ATP levels and support cell survival (Lo et al., 2023). Given these observations, we sought to test if mitochondria enable cellular recovery after USMB treatment, and if interfering with mitochondrial-related processes could heighten cell death post-USMB exposure. Collectively, we observed significantly altered mitochondrial dynamics and induction of AMPK-regulated mitophagy. Importantly, impairing mitophagy significantly boosted cell death post-USMB treatment, insinuating that co-administration of USMB with mitophagy blockers may constitute a therapeutic strategy to kill tumours.

### USMB-induced changes to mitochondrial dynamics

USMB treatment resulted in smaller but more numerous mitochondria, which is consistent with mitochondrial fragmentation rather than simply a change in morphology of individual mitochondria. Interestingly, Qiu *et al*. previously documented reduced mitochondrial area and loss of mitochondrial membrane potential following USMB treatment, aligning with our own findings (Qiu et al., 2026). Given that USMB sonoporation induces loss of ATP and metabolites and activates AMPK, we propose that AMPK promotes these USMB-driven changes to mitochondrial dynamics. Consistent with this, severe metabolic stress and mitochondrial dysfunction are known to activate AMPK, which suffices to fragment mitochondria by phosphorylating and promoting fission factors such as mitochondrial fission factor (MFF), MTFR1L, and INF2 (Chen et al., 2023; Ding et al., 2024; Ducommun et al., 2015; Herzig and Shaw, 2018; Tilokani et al., 2022; Toyama et al., 2016). Of course, other processes may alter mitochondrial morphology such as an influx of Ca^2+^ during sonoporation which can trigger fission through CaMKK-AMPK-dependent and independent mechanisms (Hatsuda et al., 2023; Hom et al., 2010; Lai et al., 2023). Ca^2+^ may also instigate an increase in oxidative stress as we detected, and this in turn, causes changes to mitochondrial dynamics (Dai et al., 2020; Przystupski et al., 2024). Smaller mitochondria may then display faster motility to redistribute mitochondria in the cell, which would be consistent with prior work showing that mitochondrial fission facilitates the redistribution and trafficking of mitochondria throughout the cell (Chan, 2020; Van Der Bliek et al., 2013). Lastly, the altered morphology and dynamics of mitochondria post-USMB may also result from mitophagosome formation, which would appear as globular rather than elongated organelles.

### USMB-mediated activation of autophagy and mitophagy

USMB treatment triggered formation of LC3 and Parkin-labelled globules, indicating that both autophagy and mitophagy are induced. Since AMPK antagonists partially abated mitophagosome numbers post-USMB exposure, this suggests that AMPK activity is a key driver of this response. Precedents for AMPK activation of mitophagy exist. For example, AMPK can stimulate autophagosome formation directly at the site of depolarized mitochondria (Liang et al., 2015). Additionally, pharmacological activation of AMPK using MK-8722 increased PINK1/Parkin-dependent mitophagy and mitolysosome formation (Longo et al., 2024) and AMPK physically interacts with phosphorylated PINK1 at Ser495 in cardiomyocytes under stress to stabilize PINK1 on the outer mitochondrial membrane leading to Parkin recruitment (Wang et al., 2018).

It remains unclear to us if USMB treatment damages mitochondria and/or if mitophagy is triggered as a scavenging process to recover building blocks. We did observe a reduction in mitochondrial membrane potential and an increase in cellular ROS, which is consistent with mitochondrial damage. In fact, others have observed that USMB sonoporation depolarizes mitochondria and/or induces ROS in tobacco BY-2 cells, HL-60, HeLa, and gastrointestinal cancer cells that may then promote apoptosis (Jia et al., 2018; Przystupski et al., 2024; Qin et al., 2013; Qiu et al., 2026; Zhong et al., 2013). However, since we also observe a rapid build-up of LC3 globules, and given the loss of proteins, ATP, and metabolites post-USMB sonoporation (Hussein et al., 2017; Lo et al., 2023), we offer that USMB-mediated stress also prompts a scavenging response to help cells survive. In fact, autophagy is often described as a nutrient-scavenging mechanism that helps maintain metabolic homeostasis in cancer cells under stress (White et al., 2021).

Consistent with the notion that mitophagy and autophagy are controlled stress responses to USMB exposure, autophagic and mitophagic flux proceed as indicated by acidification of these organelles labelled with mCherry-eGFP-LC3- and mt-Keima biosensors, respectively. In fact, USMB treatment appeared to promote acidification of autophagosomes/mitophagosomes in an AMPK-dependent manner. This is consistent with amino-acid or glucose starvation augmenting V-ATPase activity and lysosomal hyper-acidification (McGuire and Forgac, 2018; Stransky and Forgac, 2015). Moreover, AMPKα1-deficient cells retained the capacity to form LC3-positive autophagosomes under several autophagy-inducing conditions but exhibited impaired formation of acidic autolysosomes, reduced LC3-LAMP1 association, and restoration of autophagosome maturation following AMPK α1 re-expression (Jang et al., 2018). Overall, autophagy and mitophagy may serve a dual adaptive function post-USMB exposure: selectively removing dysfunctional mitochondrial material while simultaneously recycling intracellular substrates to support metabolic recovery. These functions are not mutually exclusive and may together contribute to the survival of cells that withstand the initial mechanical stressor of USMB.

### Mitophagy as a cell survival mechanism post-USMB sonoporation

USMB-induced mitophagy appears to promote cell survival as both Parkin silencing and pharmacological inhibition of PINK1 significantly stalled cell growth and/or boosted cell death following USMB (Fig. 6, 7). While BIIB-057 is reported to inhibit both PINK1 and Syk (Rasool et al., 2024), it is unlikely that Syk drives cell survival post-USMB since MDA-MB-231 cells are Syk-negative and the selective Syk inhibitor Entospletinib did not mimic BIIB-057 when used in combination with USMB (Coopman et al., 2000; Mahabeleshwar and Kundu, 2003; Wang et al., 2003). Thus, these findings support the interpretation that disruption of the PINK1-Parkin mitophagy pathway is the most likely explanation for the increased cell death; thus, the PINK1/Parkin pathway is a response to cellular damage and mediates recovery after USMB-induced stress. In cancer, the role of mitophagy and autophagy is increasingly recognized as context dependent, with evidence that it can either support tumour cell adaptation and survival or restrain tumour progression depending on tumour type, stage, and environment (Denisenko et al., 2021; Ma et al., 2020; White, 2012). This context dependence highlights the importance to further define the role of mitophagy in cellular response to USMB, particularly if mitophagy in the current context serves to recycle damaged mitochondria and/or as a scavenging process for cellular recovery.

### Limitations and conclusions of the study

Several limitations should be considered when interpreting these findings. Although observed changes in mitochondrial morphology are consistent with altered mitochondrial fission dynamics, we did not directly measure the activity of the fission machinery, nor did we delineate how USMB alters mitochondrial dynamics. In addition, our study focused specifically on PINK1-Parkin mediated mitophagy and did not investigate whether alternative forms of mitophagy may also be activated following USMB-induced stress. Nevertheless, our findings identify mitophagy as an adaptive response to USMB stress. These changes engage AMPK-associated signalling pathways resulting in the delivery of mitochondrial material to acidic compartments. Collectively, these findings suggest that pharmacological disruption of AMPK signalling or mitophagy may sensitize tumour cells to USMB sonoporation and enhance the effectiveness of USMB-based treatment strategies, which should next be tested in in vivo models such as tumour xenografts.

## Materials and Methods

### Cell Lines

MDA-MB-231 (human, female origin) and MDA-MB-468 (human, female origin) cells were obtained from American Type Culture Collection (ATCC; Manassas, VA). The MDA-MB-231 were cultured in RPMI-1640 medium supplemented with L-glutamine and sodium bicarbonate (R8758, Sigma-Aldrich; Burlington, MA). MDA-MB-468 cells were cultured in DMEM (D6429, Sigma-Aldrich). The HeLa-YFP-Parkin cell line was a gift from Dr. Peter Kim (The Hospital for Sick Children, Toronto, Canada). HeLa-YFP-Parkin cells were cultured in DMEM (D6429, Sigma-Aldrich) containing 300 μg/mL hygromycin for cell selection. All media were supplemented with 10% fetal bovine serum ([FBS]; ThermoFisher Scientific; Waltham, MA) and 1% penicillin/streptomycin (ThermoFisher Scientific). MDA-MB-231 and -468 cells were passaged every 3 to 4 days, and HeLa cells every 2-3 days to a maximum of 25 passages.

### DNA Plasmids

The pSBtet-BP-mCherry-eGFP-LC3 autophagy reporting plasmid was generated by GenScript (Piscataway, NJ). This process cloned a tandem fluorescent LC3 reporter sequence derived from ptfLC3 (#21074 Addgene; Watertown, MA) into the pSBtet-BP Sleeping Beauty transposon backbone (#60496, Addgene). ptfLC3 was originally described in (Kimura et al., 2007). In this reporter, eGFP and mCherry fluorescence mark LC3-positive autophagic structures, while the quenching of eGFP signal in acidic compartments allows for the identification of acidified LC3-positive compartments.

### Stable cell line generation

MDA-MB-231 cells were stably transfected with plasmids encoding either the doxycycline-inducible pSBtet-BP-mCherry-eGFP-LC3 autophagy reporter or mt-Keima using FuGENE HD Transfection Reagent (E2311, Promega; Fitchburg, WI), according to the manufacturer’s instructions. Twenty-four hours following transfection, cells were placed in complete RPMI-1640 medium containing 1 µg/mL puromycin (P7255, Sigma-Aldrich) for selection. Stable reporter populations were maintained under puromycin selection and passaged every 3-4 days. Reporter expression was induced using 1 µM doxycycline 24 h prior to experimentation.

### Transient siRNA transfection

For siRNA-mediated gene silencing, cells were transfected with Lipofectamine RNAiMAX (#13778075, Invitrogen; Carlsbad, CA) according to the manufacturer’s instructions. siRNAs were designed and purchased from Horizon Biodiscovery (Cambridge, UK). Cells were transfected with siRNAs for PRKN. Following 24 h of transfection, cells were incubated for 24 h in serum prior to experimentation. Knockdown efficiency was assessed by Western blotting, as described below, in HeLa cells overexpressing Parkin because endogenous Parkin abundance in MDA-MB-231 and MDA-MB-468 cells was too low to permit robust assessment of silencing by immunoblotting.

**Table 1.**
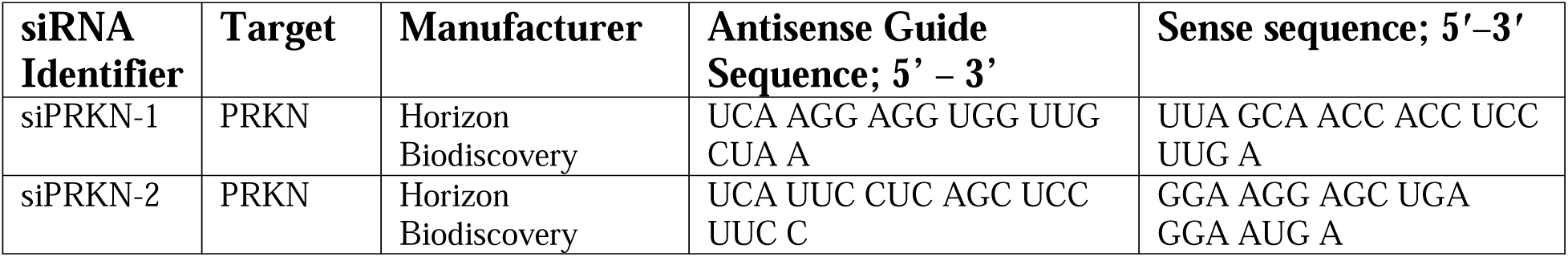
siRNA sequences used for PRKN silencing.

### Microbubble Preparation

A vial of DEFINITY microbubbles (Lantheus Medical Imaging Inc., Saint-Laurent, QC) was removed from 4°C storage and allowed to equilibrate to room temperature. After about 1 h, room temperature microbubbles were activated using a VialMix for 45 s and were used within 1 hour of activation. The vial was rolled horizontally between hands to agitate microbubbles and were then left standing upright for 30-60 s. The vial of microbubbles was punctured with an 18.5-gauge needle. Following this, 180 µL of activated microbubbles were added to 570 µL of room temperature phosphate buffered saline (PBS) both drawn using sterile syringe needles in a total volume of 750 μL.

### Ultrasound Treatment

The ultrasound setup consisted of an arbitrary waveform generator that was connected to a power amplifier (AG series Amplifier, T&C power conversion, Inc., NY, USA) that would transmit electrical signals to the ultrasound transducer. A single element flat transducer was used, with 32 mm diameter focused at 85 mm and a -6 dB beam width of 31 mm at the focal point (IL0509GP, Valpey-Fisher Inc., Hopkinton, MA, USA). Ultrasound waves were administered at 690 kPa peak negative pressure, with each sequence composed of a 32 μs pulse duration (16 cycle tone burst) and 1 kHz pulse repetition frequency, corresponding to 3.2% duty cycle. The ultrasound chamber was filled with partially degassed deionized water. The ultrasound transducer was then placed in the centre of an acoustic window of the ultrasound chamber. The acoustic window was then wiped clean with ethanol and lens paper. Before USMB exposure, wells in a six-well plate previously seeded with MDA-MB-231 cells, were filled to the brim with 16 mL RPMI-1640 media. Immediately before ultrasound exposure, microbubble/PBS mixture was shaken by hand agitation for about 1 minute. A single microbubble/PBS mixture was slowly injected into the appropriate treatment well, resulting in a concentration of 10 μL microbubbles per 1 mL of media. After injection of the microbubble/PBS mixture, the six-well dish was placed underneath the acoustic window of the ultrasound chamber. The acoustic window was lowered onto the single treatment well previously injected with the microbubble/PBS mixture. Hence, each treatment well would be exposed to ultrasound waves one at a time. It was ensured that no bubbles were formed, and a tight seal was made. The well was then exposed to an ultrasound wave at 500 kHz pulse centre frequency for 45 seconds. The administration of the microbubble/PBS mixture and the corresponding ultrasound treatment was repeated for subsequent treatment wells. As controls, each initial replicate includes an ultrasound-only treatment; if no effect, subsequent experiments simply compared no USMB and USMB exposure.

### SDS-PAGE and Western blotting

Cells were lysed using Laemmli sample buffer on ice following experimental treatments. The lysates were then collected using cell scrapers and homogenized by passing through a 27-gauge syringe 10-12 times. The lysates were subsequently heated at 95 °C, then microcentrifuged to remove insoluble material, transferring the supernatant to a new tube. Lysates were quantified for protein concentration through the usage of the Pierce Bicinchoninic Acid Assay Protein Kit (#23227, Thermo Fisher Scientific) to ensure equal protein loading before and after silencing.

Western blotting involved the use of several antibodies sourced from various retailers. The protein ladder used for the SDS-PAGE was PageRuler Plus Prestained Protein Ladder (10 to 250 kDa) (#PI26619, Thermofisher Scientific). To probe for vinculin, mouse anti-vinculin IgG (#sc-73614, Santa Cruz Biotechnology; Dallas, TX) was used. To probe for parkin, rabbit anti-Parkin primary antibody (#2132, Cell Signaling; Danvers, MA) was used. For secondary antibodies, anti-mouse IgG, HRP-linked antibody (#7076, Cell Signaling) and anti-rabbit IgG, HRP-linked antibody (#7074, Cell Signaling) were both used. Western blot PVDF membranes were imaged using the ChemiDoc Touch Imaging System by Bio-Rad. Densitometry was performed using Fiji software.

### Mitochondrial labelling

Mitochondrial labelling used MitoTracker Deep Red and MitoTracker Green (M22426 & M7514, ThermoFisher). Both were used as MitoTracker Deep Red is appropriate for live cell imaging and fixation, whereas MitoTracker Green is only appropriate for live cell imaging and is poorly retained after fixation. Both were added directly to the media for a final concentration of 100 nM, incubated for 15 min, and subsequently the media was replaced with fresh media.

### CellROX labelling

To assess intracellular ROS, MDA-MB-231 cells were incubated with 5 μM CellROX Deep Red (#C10422, Invitrogen) and imaged by live-cell spinning disk confocal microscopy. Cells were analyzed using a positive control of 5 μM sodium arsenite (NaAsO_2_) as a positive control for oxidative stress, or at 30 min and 4 h following USMB treatment. ROS levels were quantified by measuring mean fluorescence intensity per condition. ROS signal from treated cells was then compared with untreated control cells to determine treatment-dependent changes in oxidative stress.

### Pharmacological Inhibition of AMPK

To assess the contribution of AMPK activity to the cellular response following USMB sonoporation, MDA-MB-231 cells were treated with the AMPK inhibitors compound C [(dorsomorphin) # 60887S; Cell Signaling] or BAY-3827 (#2377576-35-5, Cayman Chemical; Ann Arbor, MI)(Bringas et al., 2025; Zhou et al., 2001). Compound C and BAY-3827 were used at a final concentration of 1 µM. Cells were pre-incubated with the respective inhibitors for 1 hour prior to USMB, during USMB, and 1 hour after USMB. Vehicle-treated cells received an equivalent volume of DMSO.

### Pharmacological Inhibition of PINK1

To assess the contribution of PINK1 signalling to cell survival following USMB treatment, cells were treated with BIIB-057 (#1370261-96-3, Cayman Chemical), at a concentration of 2 µM. The selective SYK inhibitor entospletinib (#T6101, TargetMol; Wellesley Hills, MA) was used at a concentration of 1 µM. Cells were pre-incubated with the respective inhibitors before USMB exposure. Vehicle-treated cells received an equivalent dose of DMSO. Cell confluence and CellTOX Green-positive dead-cell counts were monitored over 72 h using the IncuCyte system, as described below.

### Fluorescence microscopy

Fluorescence images were acquired using spinning-disk confocal microscopy. The primary microscope used was an Olympus IX83 P2ZF spinning-disk confocal microscope equipped with a UPLAPO OHR 60x 1.5 numerical aperture oil-immersion objective, a Hamamatsu ORCA-Flash 4.0 camera, and CellSens/LDI acquisition software. Additional imaging was performed on a Quorum spinning-disk system mounted on a Leica DMi8 microscope. It was operated using Quorum Wave FX and used the software MetaMorph. Images were taken using the Andor Zyla sCMOS camera. Live-cell imaging was performed using an environmental chamber maintained at 37°C and 5% CO_2_ when required. Acquisition settings were kept consistent across treatment conditions and replicates. Images were exported and analyzed as described below.

### Image Analysis

Confocal microscopy images were analyzed using Fiji/ImageJ. Mitochondrial morphological characteristics, including parameters related to mitochondrial size, shape, and network complexity were quantified using the Mitochondrial Analyzer plugin (Chaudhry et al., 2020). For time-lapse imaging experiments, mitochondrial motility was analyzed using the TrackMate plugin, which enabled detection and tracking of mitochondrial structures across sequential frames (Tinevez et al., 2017). TrackMate-derived measurements, including mitochondrial speed and displacement, were exported for statistical analysis.

Mitochondrial membrane potential was assessed by comparing TMRE fluorescence to MitoTracker Green-labelled mitochondrial area. MitoTracker Green was selected for this because it tracks all mitochondria, regardless of membrane potential. TMRE-positive mitochondrial signal was normalized to total mitochondrial area. Oxidative stress was quantified from CellROX-labelled cells by measuring the mean fluorescence intensity per cell, using identical acquisition and analysis settings across conditions within each replicate.

Autophagy was assessed in MDA-MB-231 cells expressing the tandem fluorescent mCherry-eGFP-LC3 reporter. LC3-positive puncta were identified from the mCherry channel and quantified as puncta per cell. Acidified LC3-positive structures were identified as mCherry-positive/eGFP-dim puncta, whereas non-acidified structures retained both mCherry and eGFP fluorescence. The acidified LC3 fraction was calculated as the proportion of mCherry-positive/eGFP-negative LC3 puncta relative to total puncta.

Mitochondrial association with LC3-positive structures was assessed using Coloc2 in Fiji/ImageJ. Manders’ coefficient values were calculated in order to determine the fraction of mitochondrial signal overlapping with LC3-positive puncta. Neutral and acidic mt-Keima channels were separated, and then binary masks were generated from the fluorescence signal in each channel. These masks were used to quantify fluorescence intensity in the corresponding channels, and mitophagic flux was expressed as a ratio of acidic to neutral mt-Keima signal. For all image analyses, identical methods of analysis were performed across all treatment conditions within the same experiment.

### Cell Growth and Death Analysis

MDA-MB-231 cells were analyzed for cell proliferation and death using the Incucyte SX5 Live-Cell Analysis System (Sartorius; Göttingen, DE). MDA-MB-231 cells were seeded on 6-well tissue culture plates with 2.0 x 10^5^ cells per well (83.3920, Sarstedt; Nümbrecht, DE). After approximately 12 to 16 hours of growth, cells in the USMB conditions were subjected to USMB, following the aforementioned protocol. CellTox (Promega; G8741) at a 1:10 000 dilution was added to all wells. Immediately after, 6-well plates were placed into the Incucyte apparatus for imaging. Images were acquired under 1-hour intervals under 10x magnification using brightfield phase contrast and exposure of 460 nm excitation and 524 nm emission of green fluorescence. This allowed for the excitation of the CellTox^TM^ dye, and image collection for the purposes of cell death analysis. Once imaging was completed, cells were subsequently removed from the Incucyte for disposal. Cell confluence and cell death measurements were then obtained by the Incucyte SX5 Live-Cell Analysis System.

### Statistical Analysis

Statistical analyses were performed using the software Graphpad Prism. Data are presented as mean ± standard deviation or mean ± SEM, as indicated in the figure legends. The number of independent biological replicates, cells analyzed per condition, and statistical tests used for each experiment are reported in the corresponding figure legends. For microscopy-based experiments, cell measurements of a treatment group within each independent experiment were averaged to generate a single sample-level mean. The sample means from each independent experiment were then used for statistical comparison. Assuming normally distributed data, for comparisons between two groups, experimentally paired t-tests were performed as indicated. For comparisons among three or more groups, one-way or two-way repeated measures ANOVA analyses were performed, applying the Geisser-Greenhouse correction, where applicable, followed by the recommended post-hoc tests as specified in the figure legends. If data were normalized, then non-parametric tests were employed. For IncuCyte cell death assays, area under the curve (AUC) values were calculated for each independent experimental replicate prior to statistical comparison. Differences were considered statistically significant at p < 0.05, but p values are disclosed.

## Abbreviations

ATP: Adenosine triphosphate;
AMPK: Adenosine Monophosphate-activated Protein Kinase;
AUC: Area Under Curve;
DCC: Dead Cell Count;
LC3: Microtubule-associated protein 1A/1B-light chain 3;
MFI: Mean Fluorescence Intensity;
MTDR: MitoTracker Deep Red;
MTG: MitoTracker Green;
NaAsO_2_: Sodium Arsenite;
PINK1: PTEN-Induced Kinase 1;
TMRE: Tetramethylrhodamine ethyl ester;
TNBC: Triple-Negative Breast Cancer;
USMB: Ultrasound and Microbubble Treatment

## Acknowledgements

We would like to thank Drs. Tamotsu Yoshimoro and Eric Kowarz for donating plasmids used in this work and deposited in Addgene. We thank Dr. Ella Hyatt for managing the research facilities at TMU MaRS.

## Supplemental Material

**Supplemental Figure S1. USMB sonoporation alters mitochondrial network architecture in MDA-MB-468 cells**. **A.** Representative images of mitochondria in MDA-MB-468 cells labelled with MitoTracker Green before USMB sonoporation (control), or 4 h post-USMB treatment. **B-G.** Quantification of mitochondrial morphology which includes mean area per mitochondrion (B) mitochondrial count (C) total mitochondrial area (D), mean branch length (E), mitochondrial branch junction number (F), and mitochondrial branch number (G) under untreated control conditions or 4 h following USMB treatment conditions. Data are shown as mean ± SEM from N = 4 independent experiments where 150-200 cells per condition per experiment were quantified. Statistical comparisons were performed using a two tailed Student’s t-test, and p values are as shown. Scale bar = 5 µm.

**Supplemental Figure S2. Mitochondrial network alterations following USMB sonoporation at 24 and 48 h. A.** Representative spinning-disk confocal microscopy images of MDA-MB-231 cells labelled with MitoTracker Green under untreated control conditions or at 24 h and 48 h following USMB treatment. **B-D.** Quantification of mitochondrial morphology at 24 h and 48 h following USMB treatment compared with time-matched untreated controls, including mean mitochondrial branch length (B), number of mitochondrial branch junctions (C) and number of mitochondrial branches (D). Data are presented as mean ± SEM from N = 3 independent experiments where 150-200 cells per condition per experiment were quantified. Statistical comparisons between control and USMB-treated cells at each time point were performed using a two-way ANOVA followed by Šidak’s multiple comparisons test with p-values indicated. Scale bar = 5 µm.

**Supplemental Figure S3. Parkin silencing enhances USMB-induced cell death in MDA-MB-468 cells. A.** Representative immunoblot validating Parkin silencing by the Parkin-targeting siRNA oligonucleotides. Knockdown efficiency was assessed in HeLa cells overexpressing Parkin because endogenous Parkin abundance in MDA-MB-231 and MDA-MB-468 cells was too low to permit robust assessment of silencing by immunoblotting. Vinculin was used as a loading control. **B.** Densitometric quantification of Parkin protein abundance normalized to vinculin and expressed relative to the untreated control condition. For B, statistical comparisons were performed on normalized Parkin protein abundance using Friedman’s test followed by Dunn’s multiple comparisons test. Data are presented as mean ± SD (N = 5). **C.** Incucyte images with merged phase and fluorescence images acquired over 72 h showing CellTox Green-positive cells in MDA-MB-468 cells transfected with control siRNA or siParkin, with or without USMB treatment. **D.** Quantification of cell confluence over the 72 h period, expressed as the area under the curve (AUC). **E.** Quantification of cumulative cell death over time, as expressed as the AUC of CellTOX Green-positive dead-cell counts. **F.** Quantification of cell death normalized to cell confluence, expressed as the AUC of dead-cell count/confluence. For **D-F,** statistical comparisons were performed on AUC values using a two-way ANOVA followed by Tukey’s multiple comparisons test. Data from **D-F** are presented as mean ± SEM (N=3).

**Supplemental Video S1: Mitochondrial dynamics in untreated MDA-MB-231 cells.** MDA-MB-231 cells labelled with MitoTracker Green were monitored by live-cell spinning-disk time-lapse microscopy. Single-plane images were acquired at 10 s intervals for 12 frames. Scale bar = 5 µm.

**Supplemental Video S2: Mitochondrial dynamics following USMB treatment.** MDA-MB-231 cells labelled with MitoTracker Green were monitored by live-cell spinning-disk time-lapse microscopy 4 h following USMB treatment. Single-plane images were acquired at 10 s intervals for 12 frames. Scale bar = 5 µm.

**Supplemental Video S3: Live-cell Incucyte imaging of untreated MDA-MB-231 cells.** MDA-MB-231 cells were monitored by Incucyte live-cell imaging at 1 h intervals for 72 h. Phase-contrast images were acquired alongside CellTOX Green fluorescence to visualize cell death.

**Supplementary Video S4. Live-cell Incucyte imaging of MDA-MB-231 cells transfected with non-targeting control siRNA.** MDA-MB-231 cells transfected with non-targeting control siRNA (siControl) were monitored by Incucyte live-cell imaging at 1 h intervals for 72 h. Phase-contrast images were acquired alongside CellTOX Green fluorescence to visualize cell death.

**Supplementary Video S5. Live-cell Incucyte imaging of Parkin-silenced MDA-MB-231 cells.** MDA-MB-231 cells transfected with PRKN-targeting siRNA Oligo 1 were monitored by Incucyte live-cell imaging at 1 h intervals for 72 h. Phase-contrast images were acquired alongside CellTOX Green fluorescence to visualize cell death.

**Supplementary Video S6. Live-cell Incucyte imaging of MDA-MB-231 cells following USMB treatment.** MDA-MB-231 cells were monitored following USMB treatment by Incucyte live-cell imaging at 1 h intervals for 72 h. Phase-contrast images were acquired alongside CellTOX Green fluorescence to visualize cell death.

**Supplementary Video S7. Live-cell Incucyte imaging of control siRNA-transfected MDA-MB-231 cells following USMB treatment.** MDA-MB-231 cells transfected with non-targeting control siRNA (siControl) and exposed to USMB treatment were monitored by Incucyte live-cell imaging at 1 h intervals for 72 h. Phase-contrast images were acquired alongside CellTOX Green fluorescence to visualize cell death.

**Supplementary Video S8. Live-cell Incucyte imaging of Parkin-silenced MDA-MB-231 cells following USMB treatment.** MDA-MB-231 cells transfected with PRKN-targeting siRNA Oligo 1 and exposed to USMB treatment were monitored by Incucyte live-cell imaging at 1 h intervals for 72 h. Phase-contrast images were acquired alongside CellTOX Green fluorescence to visualize cell death.

