## Supplementary figures and images for "Mitophagy is required for cell survival upon ultrasound and microbubble therapeutic exposure"

### Supplemental Figure S1

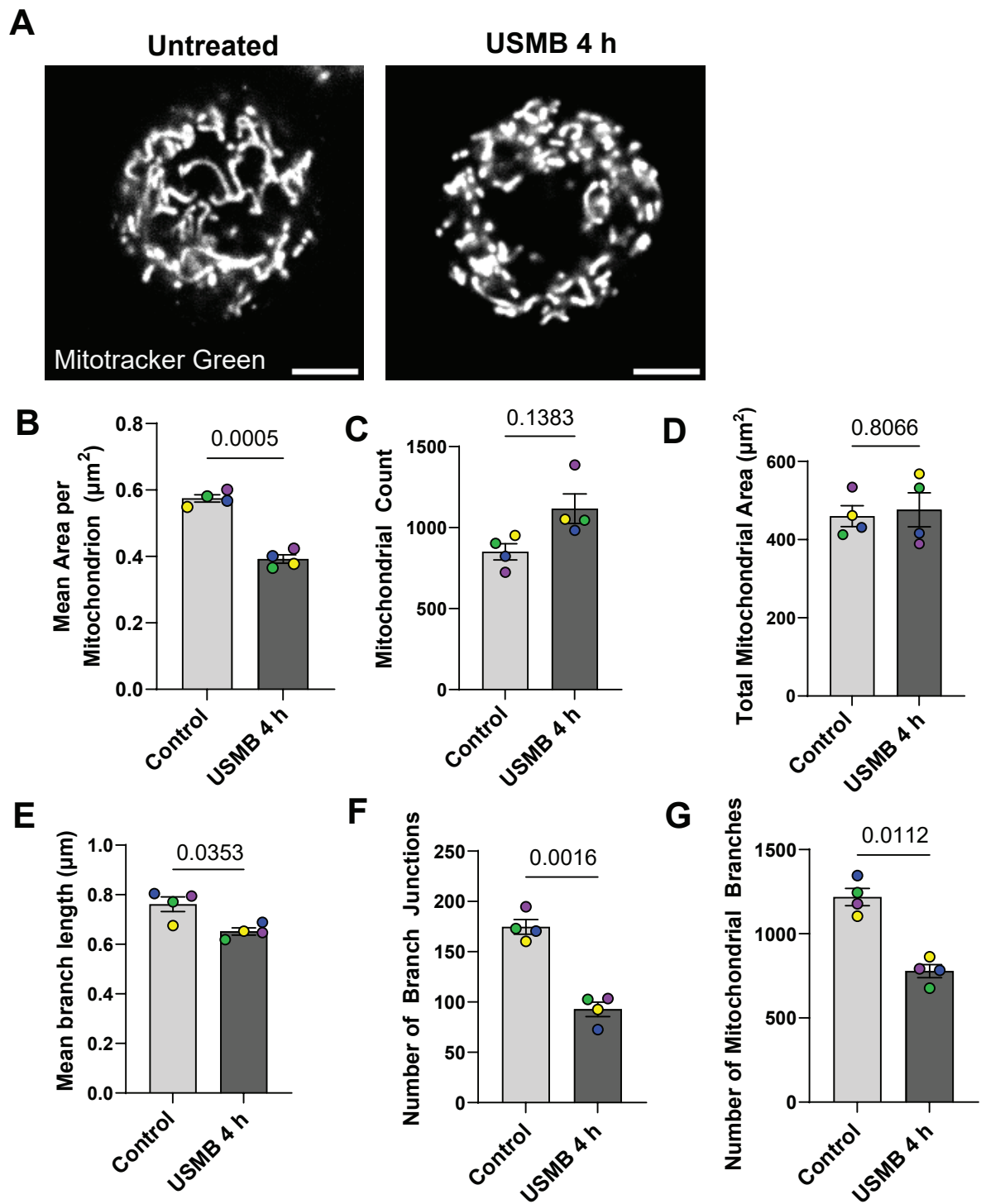

Supplemental Figure S1

### Supplemental Figure S2

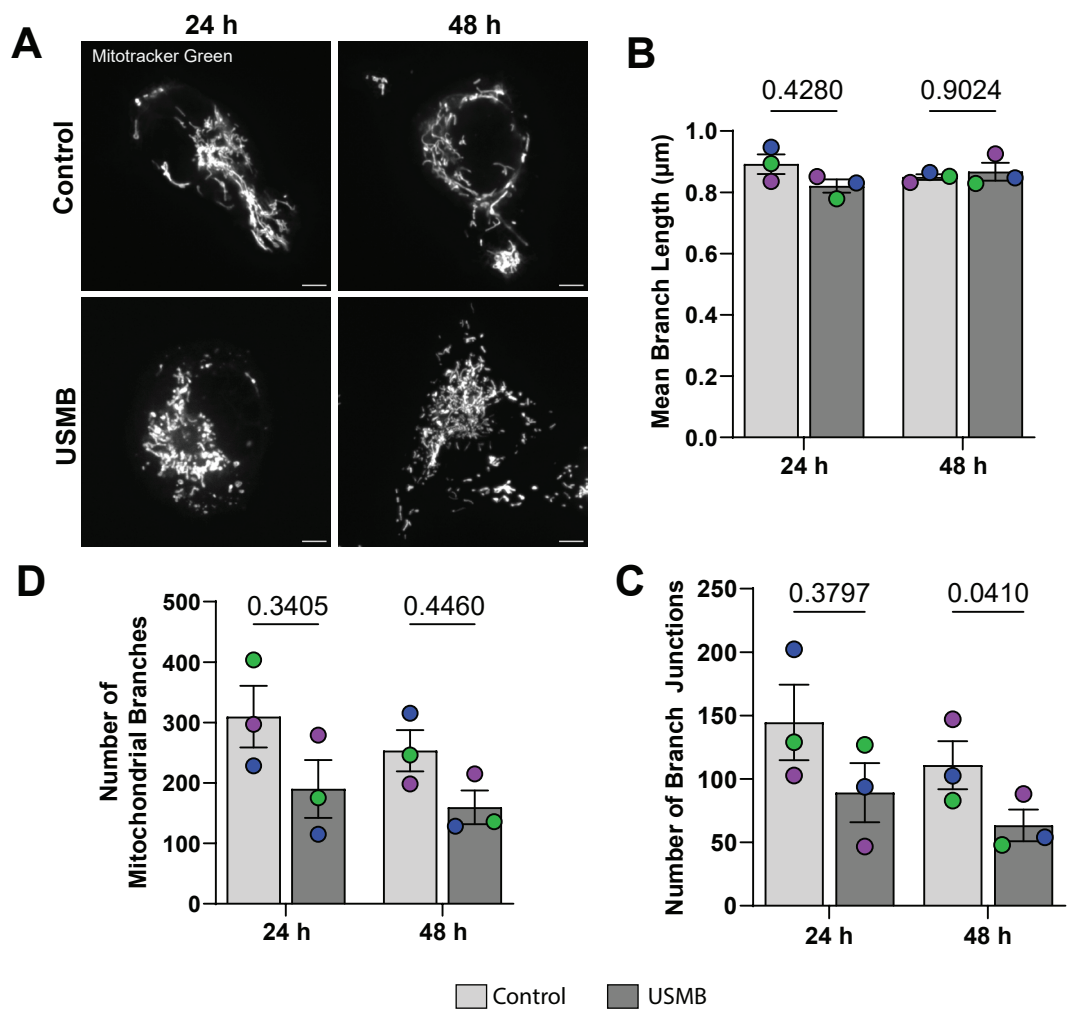

Supplemental Figure S2

### Supplemental Figure S3

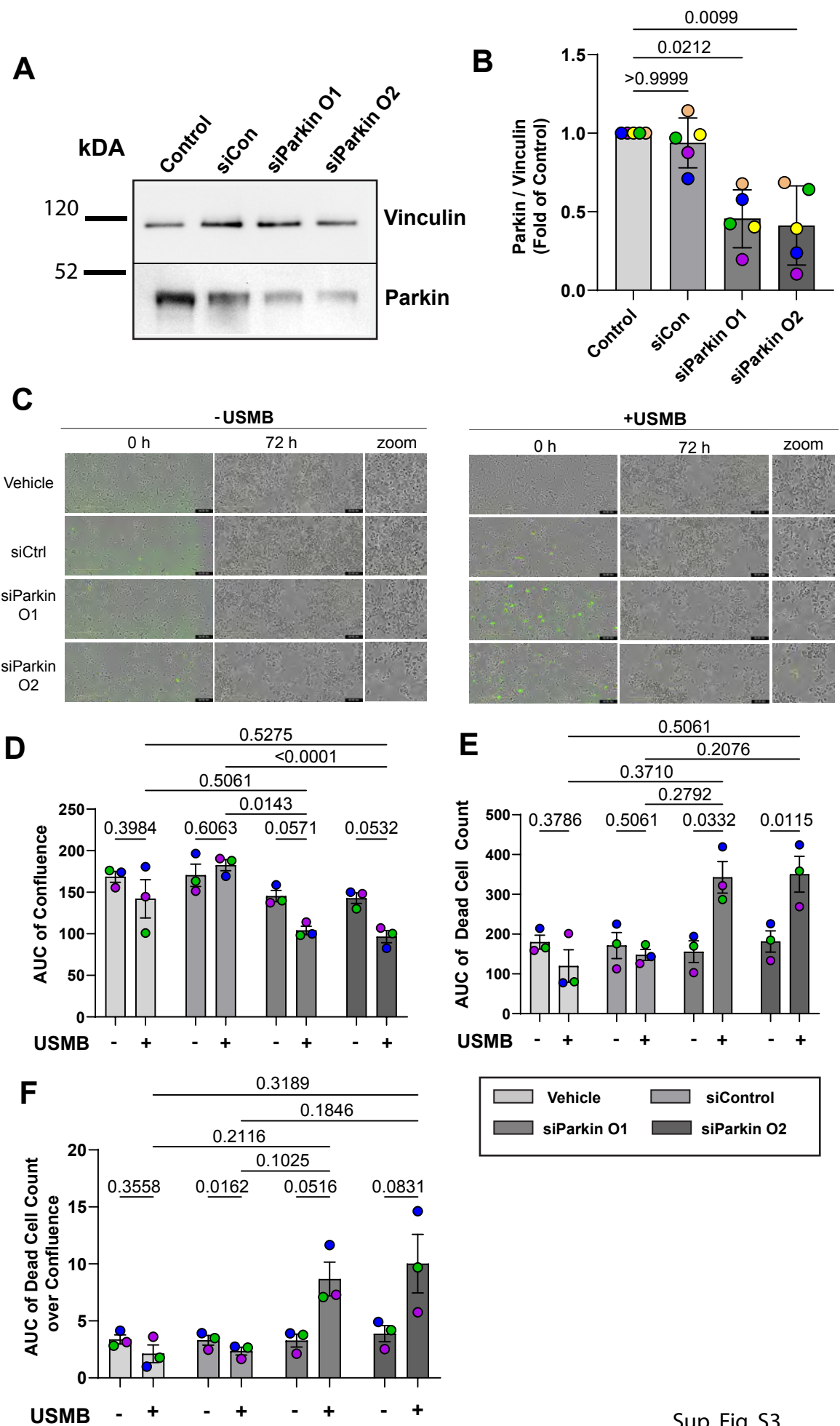

Sup. Fig. S3
